# Follow-up of a hospital cohort during the first 3,530 suspected cases of COVID-19 in Sao Jose do Rio Preto, Sao Paulo, Brazil

**DOI:** 10.1101/2021.02.04.429711

**Authors:** Carolina Colombelli Pacca, Nathalia Zini, Alice F. Versiani, Edoardo E. de O. Lobl, Bruno H. G. A. Milhim, Guilherme R. F. Campos, Marília M. Moraes, Thayza M.I.L dos Santos, Fernanda S. Dourado, Beatriz C. Marques, Leonardo C. da Rocha, Andresa L dos Santos, Gislaine C.D. da Silva, Leonardo G. P. Ruiz, Raphael Nicesio, Flávia Queiroz, Andreia F. N. Reis, Natal S. da Silva, Maurício L. Nogueira, Cássia F. Estofolete

## Abstract

**Introduction:** In a global context, COVID-19 is the most significant health threat in the present days, evidenced by the fact that, in just over four months, SARS-CoV-2 has spread to 171 countries, reaching a Pandemic status. Most patients with COVID-19 have a mild course of the disease. However, approximately 20% develop severe illness with a high mortality rate which is associated with age, comorbidities, and immunosuppression. Epidemiological studies are used to reveal the extent of viral spread in homes, communities, and hospitals. Thus, preventive and control measures can be established by the authorities.

**Objective:** In this study, patients with suspect COVID-19 symptoms who search for hospital care at the city of Sao José do Rio Preto (Sao Paulo, Brazil) were monitored, in order to identify the first case of this new disease in the region. In the first two months (March and April), more than 3000 individuals looked for the public and private health system with suspected respiratory symptoms, but only 164 (8.4%) were COVID-19 confirmed.

**Results:** From those, males (56.1%) and patients of the age distribution of 16-59 (91.2%), with diarrhea (22.2%), runny nose (25%), altered taste (15.9%), and anosmia (11.6%) presented statistical significance, although none comorbidities were related with COVID-19 occurrence. The odds ratio analysis supports this finding. Days of onset of symptoms are positively associated with whit viral load, and the same happens with the occurrence of symptoms (dyspnea and low saturation).

## 1. Background

In late 2019, Wuhan, in Hubei province, China, became the focus of the world owing to a pneumonia outbreak with unknown etiology [1]. The pathogen has been identified as novel enveloped RNA betacoronavirus (SARS-CoV-2), which has a phylogenetic similarity to SARS-CoV and that was quickly named as Coronavirus disease 2019 (COVID-19)[2]. WHO officially declared COVID-19 a pandemic on March 11, 2020 given the rapid spread of COVID-19 [3].

The first confirmed case of COVID-19 in Brazil was reported in São Paulo in February. Soon after, the virus spread through the country, and as in September 9, 2020, it had more than 4 million of cases and more than 100 thousand of deaths. The cases in the northwest of São Paulo began to be identified in the middle of March and thousands of PCRs tests have already been performed for viral diagnosis.

The SARS-CoV-2 infection mainly presents flu-like symptoms such as fever, cough and asthenia, similar to other coronaviruses. Although severe lung injury has been described at all ages, in some high-risk individuals, such as the elderly or those affected by multimorbidities, the virus is more likely to cause severe interstitial pneumonia, acute respiratory distress syndrome (ARDS) and subsequent multiorgan failure, which are responsible for severe acute respiratory failure and high death rates. Typically, affected individuals display a variable extent of dyspnea and radiological signs [8, 9]. Here, a brief description of the currently available data on the clinical features and treatment options for COVID-19.

Since the establishment of the surveillance system for COVID-19 in March until April 29, more than three thousand suspect cases were reported in the region of Sao Jose do Rio Preto. From those cases, the Laboratorio de Pesquisas em Virologia, in the Faculdade de Medicina de Sao Jose do Rio Preto, and the Central Lab from the University Hospital performed 55.1% of the required regional molecular diagnostic assays. This study describes demographic data, baseline comorbidities, clinical symptoms, presentation of clinical tests, and results of the first 1945 suspect patients who underwent RT-PCR for COVID-19 in the northwest region of São Paulo state.

## 2. Methods

This was a retrospective study of 3,530 notifications of COVID-19 and the diagnosis results from March to April 29, 2020, in Sao Jose do Rio Preto, Sao Paulo, Brazil. Real-time reverse-transcriptase polymerase chain reaction (PCR) was performed on 1,945 (55.1%) and 1,585 (44.9%) samples have not been analyzed to date. Subsequently, based on PCR results, the positive COVID-19 samples were analyzed for clinical symptoms. This study was approved by the Ethics Committee of Faculdade de Medicina de Sao Jose do Rio Preto – FAMERP (EC number 31588920.0.0000.5415).

Nasopharyngeal swabs specimens were collected from patients with suspect respiratory symptomatology who search for the public and private health system. Viral RNA was extracted using the QIAmp Viral RNA Mini Kit (QIAGEN) according to the manufacturer’s protocols. Viral RNA of SARS-COV-2 was detected by TaqMan^®^-based Real-Time PCR (Promega) using a CDC protocol with two sets of primer/probe which amplifies virus nucleocapsid (N) gene (2019-nCoV_N1 and 2019-nCoV_N2) and Human RNase P (RP) primer/probe set was included to detect the gene in control samples [4]. Samples of hospitalized patients that were originated from the University Hospital (Hospital de Base, Sao Jose do Rio Preto, Brazil) were also tested for a viral respiratory panel (Influenza A, Influenza B and Respiratory Syncytial Virus) to exclude similar pathogens and coinfections.

Standardized data were collected on demographic features, ethnicity and the presence of co-morbidities (hypertension, others cardiovascular disease, diabetes, asthma, pulmonary disease, chronic kidney disease, immunosuppression, post pregnant, neurologic disease, chromosomal disease, hematological disease, liver disease and obesity). It also collected clinical data of symptoms during the admission at the health services, including fever, cough, sore throat, dyspnea, low saturation, diarrhea, vomit, headache, myalgia, runny nose, nasal congestion, altered taste, and anosmia.

In order to measure the total of days of the symptoms, it was used the difference between the day of onset of symptoms to the day of the sample collection. This data was correlated with viral load in COVID-19 confirmed positive patients using primers N1 and N2 Ct (cycle threshold) values. It was used the software GraphPad Prism v.8 (GraphPad Software, San Diego, CA, USA) to plot graphs and to perform statistical analysis using linear regression. A paired t-test assuming Gaussian distribution was used to analyze the data. Initial symptoms and comorbidities were also correlated to the viral load. In this correlation, the median of the Ct values from both N1 and N2 primers was normalized with the internal control primer (human RNase P gene) to exclude any bias driven by low sample quality. In this case, an unpaired t-test assuming Gaussian distribution with Welch’s correction was used for data with n ≥ 20, and an unpaired t-test with Kolmogorov-Smirnov post-test was used to compare data with the non-parametric distribution. Parameters with p > 0.05 were considered to have statistical significance.

Independent variables were adopted for the final model using univariate analysis (score test). The inclusion criteria for this analysis was having statistical significance (p < 0.10) or considered significant based on clinical experience and biological plausibility [5]. Odds ratios (ORs) were adjusted with 95% of confidence interval (CI), and it was used deviance tests to ensure fit in the models, with p > 0.05 suggesting fair adjustment. Nagelkerke’s pseudo-R square was used to determine the percentage of variance explained by independent variables in the regression. The magnitude of the coefficients was assessed and used a significance level of α=5% for the evaluation of predictors in the final models. All the analyses were performed using the software SPSS, version 19 (SPSS, Inc., Chicago, IL, USA).

## 3. Results

A total of 3,530 suspected cases were reported to the survey system (Figure 1) and 1,945 samples (55,1%) were analyzed for SARS-Cov-2 by RT-PCR. 164 (8.4%) of 1,945 analysed samples by RT-PCR were confirmed for SARS-CoV-2 infection. Hospitalized patients (332/1,945; 17,06%) were tested for other respiratory viruses and it was observed circulation of Influenza A (3/332, 0.9%) and B (2/332, 0.6%) (Figure 1). The difference in days of sample colletion were compared with the day of onset of symptoms declared by the patient at the admission on health service, for negative (mean of 4.8 days) and positive (mean of 5.6 days) confirmed groups and no statiscial significance between both groups in this parameter was observed (Figure 1).

**Figure 1.**
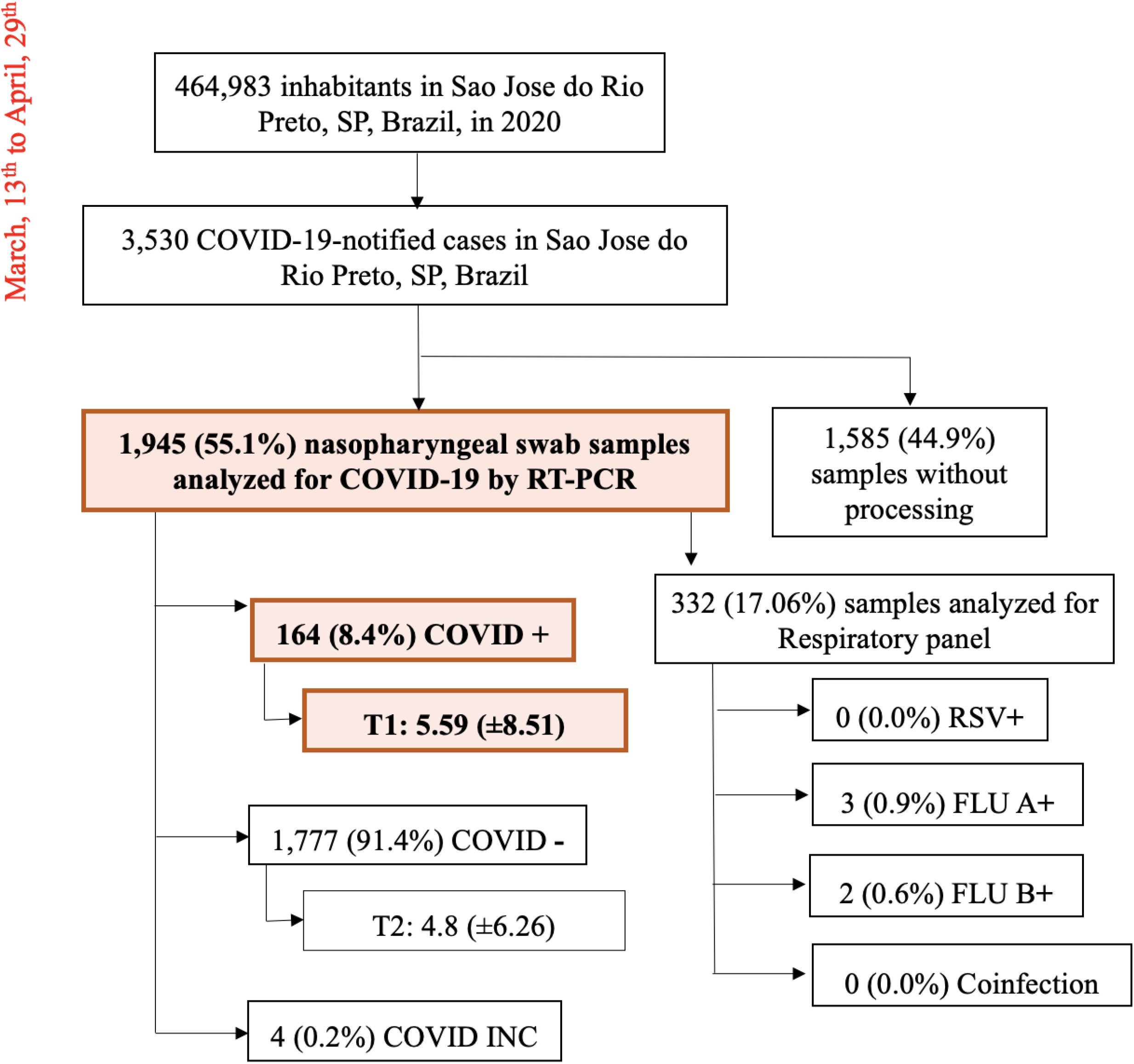
Molecular diagnostic of samples from patients with viral respiratory suspected symptoms, performed between March and April of 2020 in a hospital cohort. **Abbreviations:**RSV, Respiratory syncytial virus; INC, inconclusive

**Figure 1.**
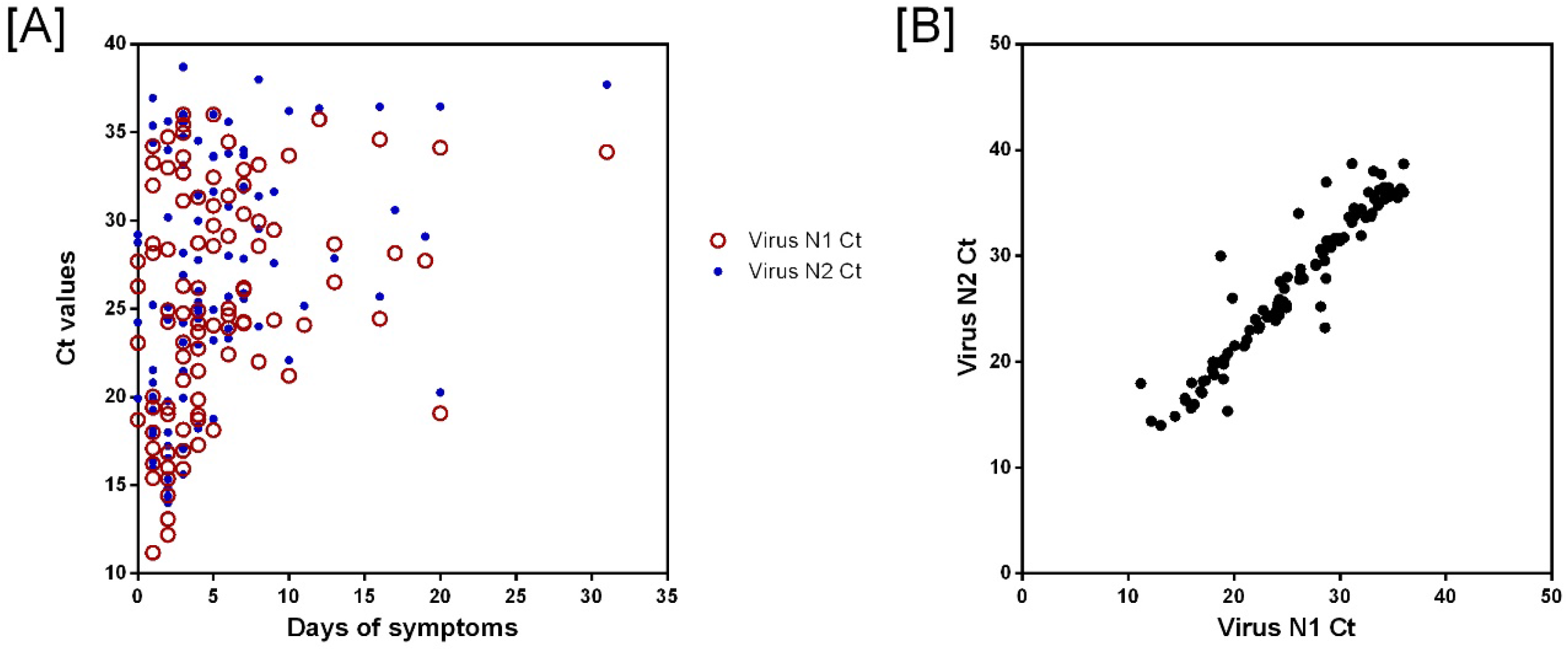
Evaluation of viral load and days of onset of symptoms. [A] Linear regression between qRT-PCR Ct values and days of onset of symptoms, p-value=0.0025. [B] Paired t-test correlation between N1 Ct value and N2 Ct value, p-value<0.0001.

The demographic and epidemiological characteristics were listed in Table 1 and 2. The median age was 40.2 years (interquartile range 22–58). About half of the positive by RT-PCR COVID patients (92/164, 56.1%) were men. The most common symptoms at illness onset were fever (770/1,480; 52%) and sore throat (700/1,483; 47.2%).

**Table 1.** Baseline characteristics of COVID-19 tested patients by RT-PCR in São José do Rio Preto, SP, Brazil, between March and April, 2020

| Demographic information | Total = 1,945 |  |  | COVID-19 positive = 164 |  |  | COVID-19 negative = 1,777 |  |  | p-value* |
| --- | --- | --- | --- | --- | --- | --- | --- | --- | --- | --- |
|  | N<br>response | N<br>positive | % or media<br>or SD | N<br>response | N<br>positive | % or media<br>or SD | N<br>response | N<br>positive | % or media<br>or SD |  |
| **Age, media** | 1,495 | 40.21 | 18 | 125 | 41.84 | 12.58 | 1,366 | 40.04 | 18.44 | 0.14 |
| **Distribution** |  |  |  |  |  |  |  |  |  |  |
| 0-15 yr | 1,495 | 98 | 6,6 | 125 | 1 | 0,8 | 1,366 | 97 | 7,1 | 0.008 |
| 16-59 yr | 1,495 | 1,211 | 81,0 | 125 | 114 | 91,2 | 1,366 | 1,093 | 80,0 |  |
| 60-79 yr | 1,495 | 140 | 9,4 | 125 | 9 | 7,2 | 1,366 | 131 | 9,6 |  |
| ≥80 yr | 1,495 | 46 | 3,1 | 125 | 1 | 0,8 | 1,366 | 45 | 3,3 |  |
| **Gender** |  |  |  |  |  |  |  |  |  |  |
| Female | 1,945 | 1,217 | 62.4 | 164 | 72 | 43,9 | 1,777 | 1,143 | 64,3 | <0.001 |
| Male | 1,945 | 728 | 37.6 | 164 | 92 | 56,1 | 1,777 | 634 | 35,7 |  |
| **Ethnicity** |  |  |  |  |  |  |  |  |  |  |
| Caucasian | 1,448 | 1,28 | 88.4 | 109 | 99 | 90,8 | 1,335 | 1,177 | 88,2 | 0.82 |
| Non-Caucasian | 1,448 | 168 | 6.5 | 109 | 10 | 9,2 | 1,335 | 158 | 11,8 | 0.51 |
| **Insurance** |  |  |  |  |  |  |  |  |  |  |
| Public | 960 | 329 | 19,4 | 74 | 22 | 29,7 | 884 | 307 | 34,7 | 0.54 |
| Private | 960 | 631 | 65,7 | 74 | 52 | 70,3 | 884 | 577 | 65,3 | 0.67 |
| **Comorbidities** |  |  |  |  |  |  |  |  |  |  |
| Hypertension | 1,318 | 51 | 3,9 | 98 | 5 | 5,1 | 1,216 | 45 | 3,7 | 0.74 |
| Others Cardiovascular disease | 1,318 | 177 | 13,4 | 98 | 16 | 16,3 | 1,216 | 160 | 13,2 | 0.58 |
| Diabetes | 1,318 | 105 | 8,0 | 98 | 13 | 13,3 | 1,216 | 92 | 7,6 | 0.28 |
| Asthma | 157 | 16 | 0,0 | 12 | 0 | 0,0 | 144 | 0 | 11,1 | - |
| Pulmonary disease | 1,318 | 74 | 5,6 | 98 | 4 | 4,1 | 1,216 | 69 | 5,7 | 0.53 |
| Chronic Kidney disease | 1,317 | 15 | 1,1 | 98 | 0 | 0,0 | 1,215 | 15 | 1,2 | - |
| Immunosuppression | 1,318 | 34 | 2,6 | 98 | 0 | 0,0 | 1,216 | 34 | 2,8 | - |
| Post pregnant | 1,317 | 4 | 0,3 | 98 | 0 | 0,0 | 1,216 | 4 | 0,3 | - |
| Neurologic Disease | 158 | 36 | 22,8 | 12 | 0 | 0,0 | 144 | 36 | 25,0 | - |
| Chromosomal disease | 536 | 2 | 0,4 | 40 | 0 | 0,0 | 494 | 2 | 0,4 | - |
| Hematological diseases | 157 | 2 | 1,3 | 12 | 0 | 0,0 | 144 | 1 | 0,7 | - |
| Liver disease | 157 | 1 | 0,6 | 12 | 0 | 0,0 | 144 | 1 | 0,7 | - |
| Obesity | 156 | 4 | 2,6 | 12 | 0 | 0,0 | 143 | 4 | 2,8 | - |
| **Healthcare Workers** |  |  |  |  |  |  |  |  |  |  |
| Yes | 777 | 423 | 54,4 | 55 | 28 | 50,9 | 722 | 394 | 54,6 | 0.70 |
| **Days of fever/symptoms** |  |  |  |  |  |  |  |  |  |  |
| Means | 1,538 | 4.59 | 4.73 | 127 | 5.06 | 5.09 | 1,411 | 4.55 | 4.69 | 0.24 |

**Table 2.**
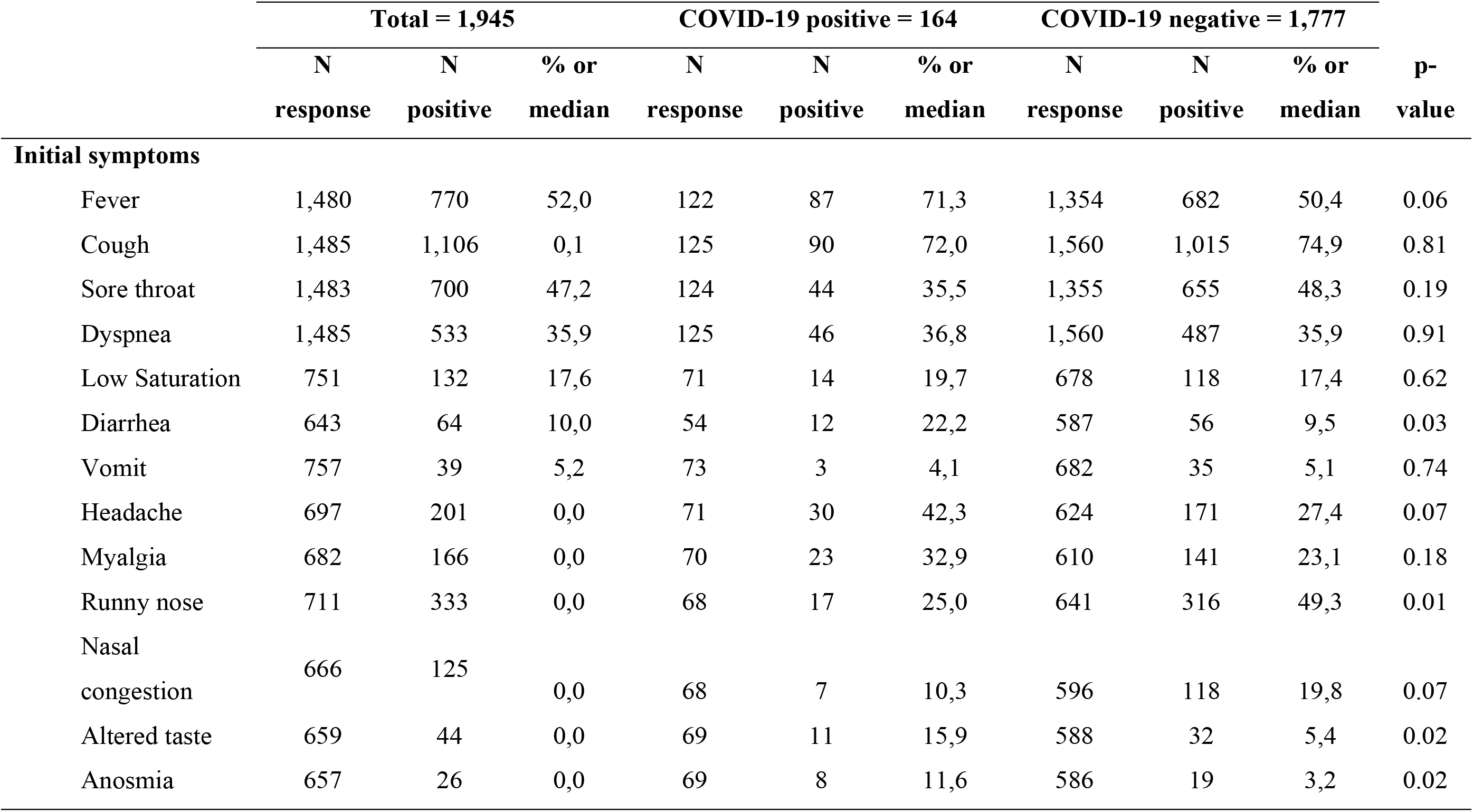
Symptoms declared by the patient at the admission at the health service, according to SARS-CoV-2 RT-PCR result.

A proportion of the infected patients had comorbidities like cardiovascular disease (16/98; 16.3%), diabetes (13/98; 13.3%), hypertension (5/98; 5.1%) and pulmonary disease (4/98; 4.1%). And a significant number of patients (28/55; 50.9%) declared themselves as a health professional, although no statistical significance was observed (Table 1). Most of the positive patients developed cough (90/125; 72%), fever (87/122; 71.3%) and headache (30/71; 42.3%) but no statistical significance was observed in these symptoms. However, diarrhea (12/54; 22.2%), runny nose (17/68, 25%), altered taste (11/69; 15.9%) and anosmia (8/69; 11.6%) presented statistical significance (Table 2).

To compare the symptoms or demographic data between those who had COVID-19 confirmed diagnostic with those who had not, it was performed a multivariate analysis. In addition to clinical experience and biological plausibility, it was used univariate analysis to select variables for the final model (p<0.10). It was observed an OR>1 in the parameters: age of 16-59, gender (masculine), fever, dyspnea, anosmia, altered taste, and headache (Table 3). Although many parameters achieve the OR>1 the ones that the 95% CI overlap the null value (OR=1) were disregarded (Table 3)

**Table 3.** Univariate and multivariate analysis (odds ratio), according to demographic characteristics and symptomatology in the disease onset

| Variables | Univariate analysis<sup>†</sup> |  | Multivariate analysis<sup>‡</sup> |  |
| --- | --- | --- | --- | --- |
|  | OR | 95% CI | OR | 95% CI |
| Age |  |  |  |  |
| (ref: 0-15 yrs) |  |  |  |  |
| 16-59 yrs | 7.78 | 1.35-70.84 | 7.87 | 1.02-60.69 |
| 60-79yrs | 8.26 | 1.06-64.60 | 4.12 | 0.46-37.01 |
| > 80 yrs | 1.73 | 0.11-28.17 | 2.71 | 0.16-47.40 |
| Gender (masc) | 2.36 | 1.71-3.26 | 4.72 | 2.60-8.59 |
| Ethnicity (Caucasian) ¶ | 1.68 | 0.68-2.60 | - | - |
| Fever | 2.45 | 2.63-3.68 | 2.07 | 1.14-3.76 |
| Cough ¶ | 0.86 | 0.57-1.30 | - | - |
| Odynophagia | 0.59 | 0.40-0.86 | 0.64 | 0.35-1.16 |
| Dyspnea ¶ | 1.04 | 1.71-1.52 | - | - |
| Respiratory discomfort ¶ | 0.57 | 0.25-1.27 | - | - |
| Desaturation ¶ | 1.17 | 0.63-2.16 | - | - |
| Diarrhea ¶ | 1.65 | 0.74-3.67 | - | - |
| Vomiting ¶ | 0.79 | 0.24-2.64 | - | - |
| Anosmia | 3.91 | 1.64-0.32 | 2.39 | 0.67-8.54 |
| Altered taste | 3.30 | 1.58-6.88 | 2.22 | 0.78-6.31 |
| Myalgia | 1.63 | 0.96-2.77 | 1.48 | 0.78-2.78 |
| Runny nose | 0.34 | 0.19-0.61 | 0.65 | 0.34-1.24 |
| Nasal congestion | 0.46 | 0.21-1.04 | 0.90 | 0.35-2.29 |
| Headache | 1.94 | 1.17-3.20 | 1.57 | 0.86-2.88 |

† Along with the clinical experience and biological plausibility, it was used univariate analysis to select variables for the final model ( $p < 0.10$ ). ‡ Deviance test: $P > 0.05$ . § Nagelkerke's pseudo-R square = 0.20. ¶ Ethnicity, cough, dyspnea, respiratory discomfort, desaturation, diarrhea, vomiting had a $p$ -value $> 0.10$ by Score test ( $p = 0.51$ ; $p = 0.48$ , $p = 0.84$ , $p = 0.16$ , $p = 0.63$ , $p = 0.22$ , $p = 0.70$ respectively). CI: confidence interval. OR: odds ratio.

In Figure 2, the Ct values of N1 and N2 target were compared with the days of onset of symptoms for each positive sample analyzed. It was observed lower Ct values in patients in which the sample were collected in the first five days of symptoms (Figure 2A). However, one patient with declared initial symptoms of over 20 days also presented Ct values below 20. It is important to note that the Ct values between N1 and N2 show a positive linear correlation and a p-value <0.0001 in a paired t-test (Figure 2B).

**Figure 2.**
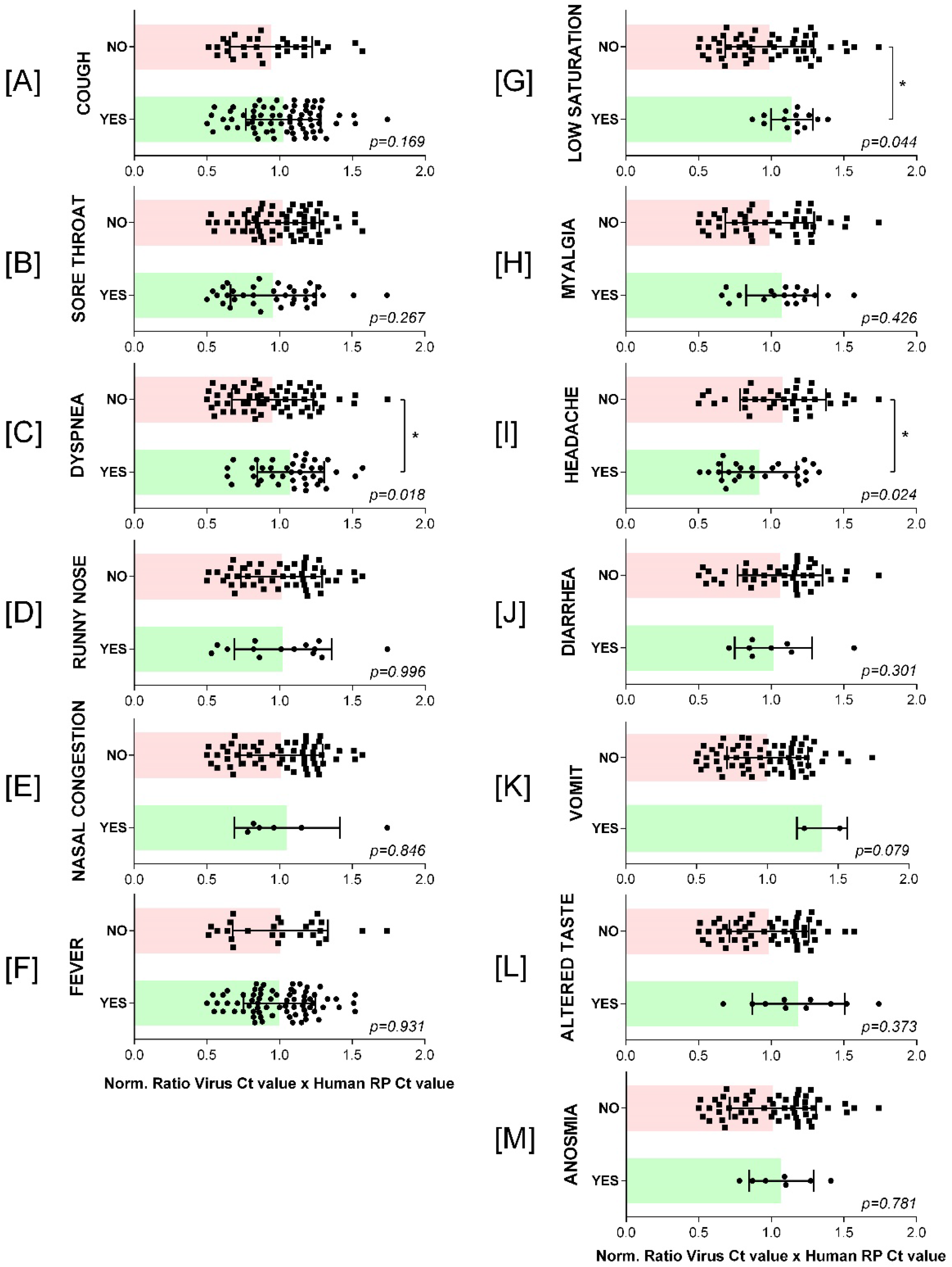
Evaluation of the correlation between viral load (normalized Ct values) and the occurrence of symptoms in patients’ COVID confirmed positives. [A] Cough, [B] Sore throat, [C] Dyspnea, [D] Runny nose, [E] Nasal congestion, [F] Fever, [G] Low saturation, [H] Myalgia, [I] Headache, [J] Diarrhea, [K] Vomit, [L] Altered taste, [M] Anosmia. Green bars: the presence of symptoms. Red bar: the absence of symptoms.

To evaluate if a higher viral load is associated with the occurrence of symptoms, it was compared Ct values (primers N1 and N2) with the symptoms and normalized with the internal control primer (human RNase P gene). Figure 3 shows that Dyspnea and Low saturation are positively related to the occurrence of COVID-19 (Figure 3C and 3G). However, headache is negatively associated with the event of the disease. Although this pattern maintains when we analyze the N1 primer alone (Figure Suppl. 1). Headache does not have statistical significance when analyzed by the N2 Ct value (Figure Suppl. 2).

## 4. Discussion

The first confirmed COVID-19 case in Sao Jose do Rio Preto region occurred on March 13, 2020. In this day, Brazil count only 34 confirmed cases, which means that this city was one of the first ones in the national territory to identify the transmission of the virus and the city was able to propose effective quarantine and lockdown measures that allowed for a slower spread of the virus. In a collaboration between the Public Health Office of Sao Jose do Rio Preto and the local Faculdade de Medicina (FAMERP), facilities from the Laboratorio de Pesquisas em Virologia were designed to perform the molecular diagnosis of COVID-19 as soon as the first case was declared at the Sao Paulo state. An emergency system was implemented, with rapid collection of samples, processing and diagnostic tests.

In the first two months of this operation, 3,530 suspected cases were reported to the health system (Figure 1), 55.9% of the subjects had biological samples collected following clinical criteria and analyzed by qRT-PCR (98.5%) or rapid test (1.5%). Only 164 of 1,945 qRT-PCR samples (8.4%) were confirmed positive for COVID-19 (Tabel 1). Considering the rapid test assay, these numbers increase to 178 (9%) of SARS-CoV-2 occurences in Sao Jose do Rio Preto (Figure 1). To ensure a proper diagnostic of patients who required hospitalization (N=332), a respiratory diagnostic panel for Influenza virus (A/B) and Respiratory syncytial virus (RSV) was performed and only Influenza A (0.9%) and B (0.6%) were detected (Table 1).

Clinical symptoms and comorbidities have been observed since the beginning of the pandemic. The results showed that the biggest number of positive cases were observed among patients median aged 40.2 years (interquartile range, 22–58 years) and the majoroty of them were male (56.1%) (Table 2). These data corroborate Souza and collaborates, who have shown that the median age of the first patients diagnosed with COVID-19 in Brazil was lower than that observed in other countries [6, 7].

Most symptoms reported by patients with COVID-19 were cough (72.0%), fever (71.3%) and headache (42.3%) and the comorbidities found were cardiovascular disease (16.3%), diabetes (13.3%), hypertension (5.1%) and pulmonary disease (4.1%) but no statistical significance was observed among these symptoms. These symptoms were also observed in other studies at the beginning of the pandemic [2, 8, 9]. The symptoms that presented a p-value lower than 0.05 are diarrhea, runny nose, altered taste, and anosmia (Table 3). However, some signs (as headache and nasal congestion) presented a tendency to be statistical significant if the number of positive confirmed patients were more significant.[10]

Although many reports correlate key comorbidities, such as cardiovascular disease and diabetes, with severity in the clinical progression of COVID-19, this cohort is an open study that did not split the patients by the outcome (mild, severe, death). This fact could explain why any of the evaluated comorbidities present statistical significance between confirmed positive and negative patients. The same analysis is pertinent to the healthcare workers parameter (Table 2).

To compare a symptom or demographic data between those who had COVID-19 confirmed diagnostic or not, it was performed a multivariate analysis. In addition to clinical experience and biological plausibility, it was used univariate analysis to select variables for our final model (p<0.10) and it was observed significant difference in age of 16-59, gender (male), fever, dyspnea, anosmia, altered taste, and headache (Table 4). It was observed the same characteristics in other studies during the pandemic caused by SARS-CoV-2 [11–13].

Some studies have found that the peak load of SARS-CoV-2 probably means that a higher viral load means lower Ct value, which is related to more viruses present at the biological sample. In Figure 2, the Ct values of primers N1 and N2 are compared with the days of onset of symptoms for each positive patient analyzed. As expected, lower Ct values are observed in patients in which the sample was collected in the first five days of symptoms (Figure 2A). However, one patient who has declared initial symptoms over 20 days also presented Ct values below 20. It is important to note that the Ct values between N1 and N2 show a positive linear correlation and a p-value <0.0001 in a paired t-test (Figure 2B). [14, 15]

The final goal in this study was to evaluate if a higher viral load is associated with the occurrence of certain symptoms. The CDC qRT-PCR protocol has used the protein of the viral nucleocapsid (N) as the viral target. In this protocol, two regions (N1 and N2) of this target protein must be detected to confirm the SARS-CoV-2 presence. To this, it was collected both Ct values (primers N1 and N2) and normalized with the internal control primer (human RNase P gene), considering that the internal control is related to the quality of the sample collection. Figure 3 shows that dyspnea and low saturation are positively related to the occurrence of COVID-19 (Figure 3C and 3G). However, headache is negatively associated with the event of the disease. Although this pattern maintains when the N1 primer was analyzed alone (Figure Suppl. 1), headache does not have statistical significance when analyzed by the N2 Ct value (Figure Suppl. 2).

## Acknowledgment

We dedicate this case report to our fellow frontline colleagues who are facing the challenges of COVID-19 response. We acknowledge the organization founder this research. MLN is supported by the São Paulo Research Foundation (FAPESP)’s COVID Program (Grant No. 2020/04836-0 to MLN) and is a Brazilian National Council for Scientific and Technological Development (CNPQ) Research Fellow. This work is supported by CAPES (Grant #001). AFV is supported by a FAPESP Fellow grant (#18/17647-0). GRFC is supported by a FAPESP Fellow grant (#20/07419-0). BCM is supported by a FAPESP Scholarship (#19/21711-9). BHGAM is supported by a FAPESP Scholarship (#19/06572-2). The funders had no role in the design of the study, collection, analyses, or interpretation of data, writing of the manuscript, or in the decision to publish the results.

## Ethics Approval

This study was approved by the FAMERP Ethical Review Board (EC number 31588920.0.0000.5415).

## Author contributions

**Carolina Colombelli Pacca**: Conceptualization, Methodology, Validation, Writing - Original Draft. **Nathalia Zini**: Conceptualization, Data Curation, formal analysis. **Alice F. Versiani**: Conceptualization, Data Curation, **Edoardo E. de O. Lobl:** Review & Editing. **Bruno H. G. A. Milhim**: Data Curation. **Guilherme R. F. Campos**: Data Curation. **Marília M. Moraes**: Data Curation. **Thayza M.I.L dos Santos**: Data Curation. **Fernanda S. Dourado**: Data Curation. **Beatriz C. Marques**: Data Curation. **Leonardo C. da Rocha**: Data Curation. **Andresa L dos Santos**: Data Curation. **Gislaine C.D. da Silva**: Project administration, Resources. **Leonardo G. P. Ruiz**: Data Curation. **Raphael Nicesio**: Data Curation**. Flávia Queiroz**: Data Curation. **Andreia F. N. Reis**: Data Curation**. Natal S. da Silva**: Data Curation, formal analysis. **Maurício L. Nogueira**: Funding acquisition, Supervision. **Cássia F. Estofolete**: Supervision, Writing - Review & Editing

## Supporting Information

**S1 Figure.**
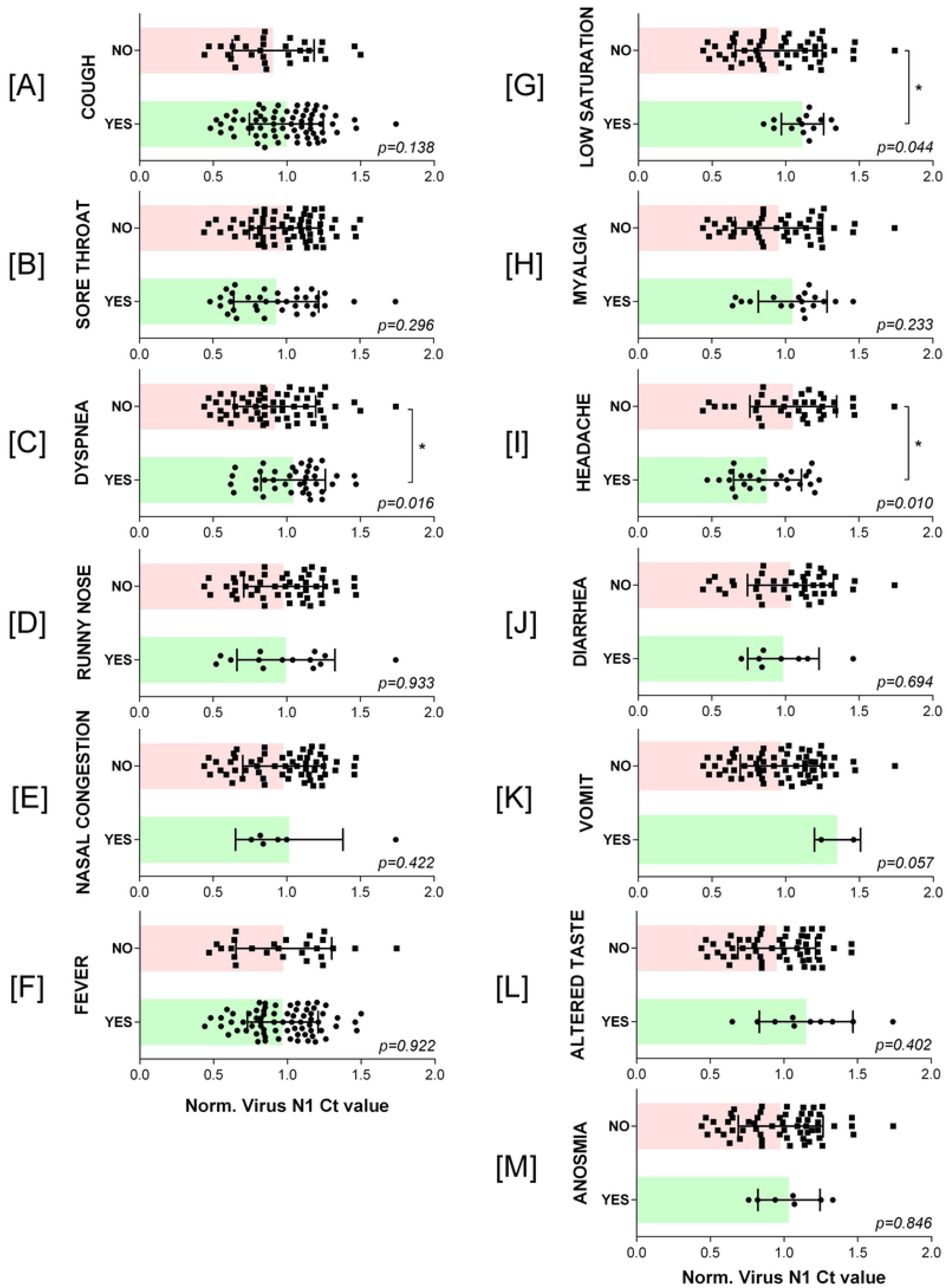
Evaluation of the correlation between viral load according N1 Ct values analyzed alone and the occurrence of symptoms in patients’ COVID confirmed positives.

**S2 Figure.**
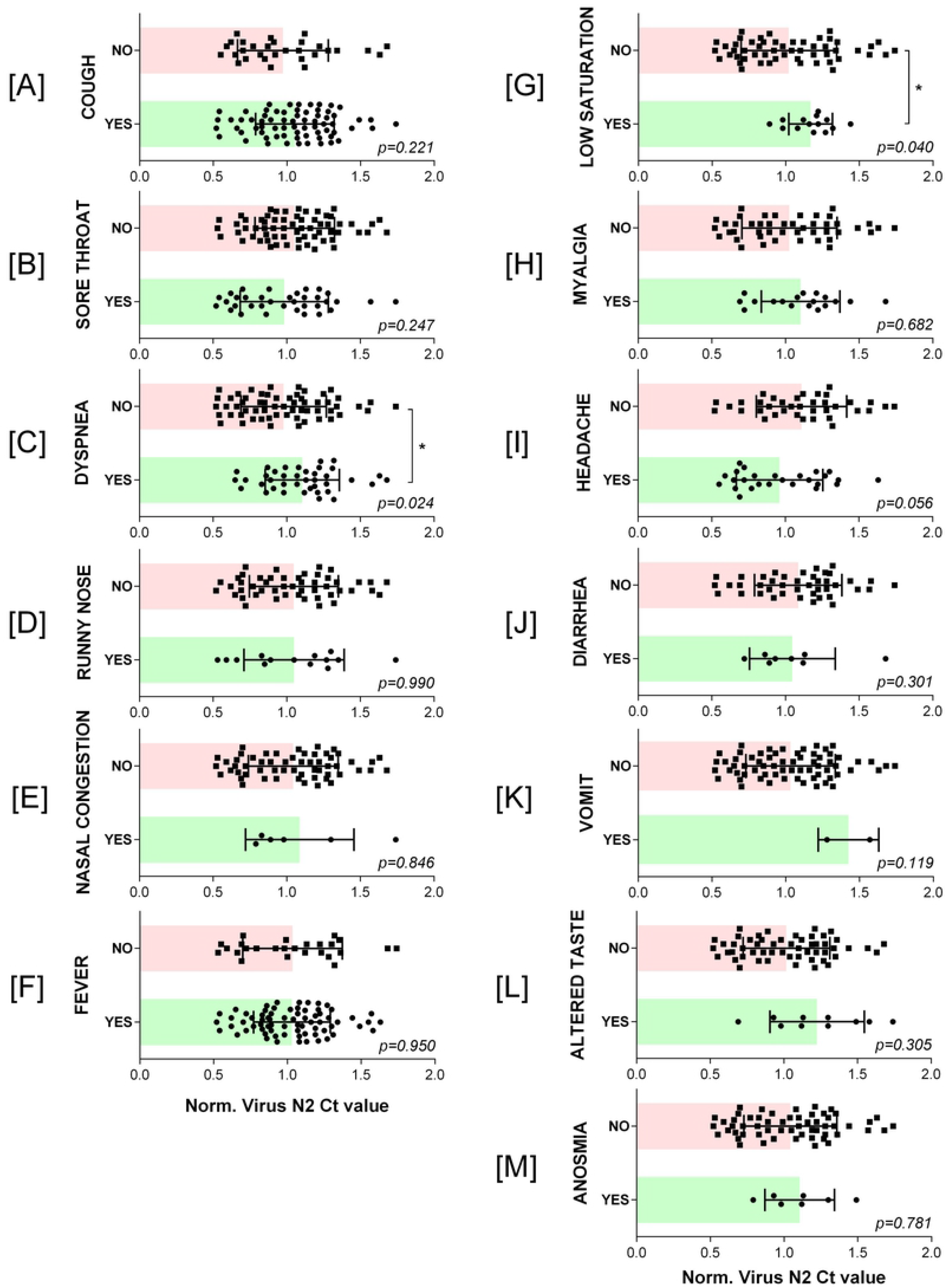
Evaluation of the correlation between viral load according N2 Ct values analyzed alone and the occurrence of symptoms in patients’ COVID confirmed positives.

